# A Machine Learning Approach to the Recognition of Brazilian Atlantic Forest Parrot Species

**DOI:** 10.1101/2019.12.24.888180

**Authors:** Bruno Tavares Padovese, Linilson Rodrigues Padovese

**Affiliations:** Bunin Tech, São Paulo, Brazil; Polytechnic School, University of São Paulo (USP), São Paulo, Brazil

**Keywords:** machine learning, neural networks, bird detection, Random Forest, MFCC

## Abstract

Avian survey is a time-consuming and challenging task, often being conducted in remote and sometimes inhospitable locations. In this context, the development of automated acoustic landscape monitoring systems for bird survey is essential. We conducted a comparative study between two machine learning methods for the detection and identification of 2 endangered Brazilian bird species from the Psittacidae species, the *Amazona brasiliensis* and the *Amazona vinacea*. Specifically, we focus on the identification of these 2 species in an acoustic landscape where similar vocalizations from other Psittacidae species are present. A 3-step approach is presented, composed of signal segmentation and filtering, feature extraction, and classification. In the feature extraction step, the Mel-Frequency Cepstrum Coefficients features were extract and fed to the Random Forest Algorithm and the Multilayer Perceptron for training and classifying acoustic samples. The experiments showed promising results, particularly for the Random Forest algorithm, achieving accuracy of up to 99%. Using a combination of signal segmentation and filtering before the feature extraction steps greatly increased experimental results. Additionally, the results show that the proposed approach is robust and flexible to be adopted in passive acoustic monitoring systems.

## 1. Introduction

Recognition of animal species by their vocalization has been employed for several years for identification of animals and localization of individuals. Governments, Non-Governmental Organizations (NGO) and researchers invest huge amounts of resources in applications such as conservation initiatives, environmental monitoring and research studies (Stephens et al., 2016; Simmonds and Isaac, 2007; Rosenzweig and Parry., 2004; Edenhofer., 2015). Particularly concerning bird recognition in tropical forests of Brazil, identification by their vocalizations is the only viable option as visibility in these types of landscapes is very limited (Retamoza et al., 2018), even at daytime. However, although there have been considerable advances in monitoring technology and artificial intelligence techniques for automatic classification, manual surveys are still widely used (Priyadarshani et al., 2018). These manual implementations of acoustic monitoring and identification often impose a series of problems such as being time-consuming, costly, manpower heavy, logistically complicated and require extensive training of a human operator to correctly identify the sounds while still being error prone (Sebastián‐González, 2015; Bardeli et al., 2010).

Notably, the identification of endangered bird species and their population size is of deep interest to the scientific community and NGOs interested in evaluating ecological conservation. Usually having very low population density, efforts towards correct monitoring encompasses large areas, requiring even further resources. Furthermore, when different species from the same family are present in the same ecosystems, correctly identifying vocalizations is a very difficult task even for a trained human operator.

Passive acoustic monitoring (PAM) systems have supported researchers by providing an efficient way to monitor large remote areas with an array of unmanned equipment capable of months to even years of uninterrupted acoustic data collection (Blumstein, 2011; Johnson et al., 2002). These systems can be designed for a wide field of applications, for instance, biodiversity assessment, surveillance of conservation units, population census surveys and pest detection (Chesmore, 2004). Furthermore, recent developments have made these systems low-cost, accessible, simple to operate and requiring little maintenance (Priyadarshani et al., 2018). On the other hand, long times of uninterrupted acoustic data makes manual analysis of events impractical without tools to automatically identify and classify acoustic events (Sebastián‐González, 2015).

Rapid advances in the field of pattern recognition and machine learning are leading to the development of tools and systems capable of analyzing huge datasets in several fields in unprecedented ways (Yegnanarayana, 2009; Cortes and Vapnik, 1995). Therefore, the implementation of machine learning models to analyze and classify acoustic signals from large datasets is natural. While there has been some research conducted in acoustic species identification, it is still a peripheral field of pattern recognition and signal processing, and most studies are still devoted to the problem of speaker identification and analysis (Le-Qing, 2011). Additionally, the numbers of studies in the field are insufficient to meet the challenge of animal recognition due to the amount of biodiversity existent across numerous landscapes and to the increasing number of environmental issues that needs to be tackled.

In this context, the incorporation of machine learning models alongside PAM systems enable the development of fully automated monitoring and recognition systems capable of unmanned long-term monitoring in inhospitable areas (Chesmore, 2004). Furthermore, these solutions need to be both robust and flexible in order to accommodate the diverse landscapes and ecosystems such as tropical forests, aquatic systems and for monitoring nocturnal species activity (Miles et al., 2006; Depraetere et al., 2012; Berg et al., 2006; Johnson et al., 2002).

Among the studies conducted for animal detection and classification, machine learning has usually been employed to identify species that have distinct vocalizations that are usually loud and clear such as insects, amphibians, birds and whales, to name a few. For example, (Brandes, 2008) exposes a series of commonly found problems in bird monitoring activities, as well as motivation towards PAM systems for avian survey. Additionally, the author discusses several hardware approaches towards the development of these systems, such as possible microphones for automated recordings, hardware for scheduling recordings, embedded systems, and even smartphones. Finally, an overview of the prominent techniques used in software, to automatically classify the collected recordings, is described. In it, Machine learning techniques are considered along with commonly used feature extraction methods for acoustic analysis.

Concerning amphibians, (Jaffar and Ramli, 2013) study the automatic identification of 10 frog species vocalizations by using machine learning and an automatic syllable segmentation procedure to extract features. Firstly, the recordings were processed by the proposed syllable segmentation method into a set of syllables. Next, each syllable is subjected to pre-emphasizing, framing and windowing. With the pre-processed syllables, a feature extraction step is conducted with the Mel-Frequency Cepstral Coefficients (MFCC) and Linear Predictive Coding (LPC) algorithms. Finally, the classic *kNN* classifier is used to train the model.

As for insects, (Le-Qing, 2011) aims to provide a method to recognize insects to aid in pest management and control. The method consists of a pre-processing stage composed of signal normalization, pre-emphasis to boost high frequency components and signal segmentation into a series of samples. Then, features are extracted using MFCC and a Probabilistic Neural Network (PNN) is trained with the resulting features. The results showed recognition rate of up to 96% from 50 different insect vocalizations.

In another work, (Johnson et al., 2002) discuss the special difficulties found particularly in nocturnal bird survey (i.e. owls). The authors propose an automated monitoring technique, using specialized tools for optimizing both time and resources during nocturnal surveys and present the results of field tests when applying their method. Finally, a series of further issues that needs to be considered are discussed, such as hardware failure in a remote location.

In (Pace, 2008), humpback whale song classification is studied, and a combination of several feature extraction and machine learning methods are compared. Specifically, the MFCC, LPC and real cepstrum coefficients features were combined with k-means, a clustering algorithm for classification, and the Multilayer Perceptron (MLP), and Artificial Neural Network (ANN).

In its turn, (Lopes et al., 2011) focus on the automatic identification of a large number of bird species from the Southern Atlantic Brazilian Coast. Firstly, a series of pre-processing steps are conducted to ensure the quality of the recordings. Next, features are extracted from each bird segment using the MARSYAS framework and the *kNN*, MLP, Naïve Bayes and SVM methods are trained in order to classify the samples.

An interesting method proposed by (Selin et al., 2006) use a wavelet approach to recognize inharmonic and transient bird sounds. After a usual pre-processing step of noise reduction and signal segmentation, the approach consists of using wavelet decomposition as the feature extraction step. With it, four features were extracted and fed to two types of Neural Networks, the unsupervised self-organizing map (SOM) and the MLP achieving results of up to 96% of accuracy.

In what concerns automatic bird monitoring and classification, the literature describes two major challenges still present today: the difficulty to access remote locations in order to conduct bird survey, and the ability to effectively process huge datasets with multiple acoustic events. These challenges are usually overcome by using remote sensors, to automatically collect vocalizations, or by an automatic classification approach to detect and classify the collected data, or a combination of both. Furthermore, due to its widespread use in speaker identification problems and very well documented results, almost all studies use the classic MFCC method for feature extraction. Regarding machine learning methods, although several classical methods were tested, their focus was applied in pre-processing stages by enhancing the recordings quality and segmenting the signal, or during the feature extraction step.

All in all, this work aims to create an automated recognition and detection method for the identification of two endangered Brazilian bird species, the *Amazona brasiliensis* and the *Amazona vinacea* from the Psittacidae family. Additionally, 5 other bird species from the Psittacidae family are considered in this study as to verify the methods capability to handle the difficult task described in the previous paragraph. For this purpose, we conducted a comparative study between 2 machine learning methods using MFCC features for both, and a series of pre-processing steps.

The red-tailed amazon (*Amazona brasiliensis*), is a species of parrot from the Psittacidae family, and is endemic to the Atlantic Forest in Brazil. This species, that lives in forests and mangroves near the coast, had around 2000 estimated individuals left in 1991 due to loss of habitat and illegal poaching (BirdLife International, 2017). Due to recent conservation initiatives, its population is increasing and has recovered to about 6000 individuals and is now considered as “near threatened” by the IUCN (International Union for Conservation of Nature) in its Red List of Threatened Species (BirdLife International, 2017). However, illegal poaching is still considered a major risk as this species is a prime target due to its vibrant color.

The vinaceous-breasted amazon (*Amazona vinacea*) is another parrot species from the Psittacidae family and can be found in tropical and subtropical regions of Brazil, Argentina and Paraguay. With less than 3000 individuals remaining and with its population spread out in a large area, it is classified as “endangered” by the IUCN (BirdLife International, 2017).

In summary, this paper describes a method for the detection and classification of two endangered Brazilian bird species. This method is also capable of being easily implemented in systems that are deployed in remote locations. Furthermore, this paper aids researchers and NGOs in handling the difficult task of differentiating a particular species among several other similar acoustic vocalizations.

## 2. Materials and Methods

### 2.1. Overview

Concerning the objective of identifying the *Amazona brasiliensis* and *Amazona vinacea*, we compared two different machine learning models which are the Random Forest algorithm and an artificial neural network (ANN), the multilayer perceptron (MLP). To achieve this, we adopted the following methodology, illustrated in Figure 1. First, the collected recordings are run through a pre-processing step composed of signal segmentation and filtering the desired band. Next, MFCC features are extracted and normalized (only for the Neural Network) aiming at capturing relevant information to the identification problem. Finally, in the last step, the resulting feature vector is fed to the chosen classification algorithms and we compute the results.

**Figure 1.**
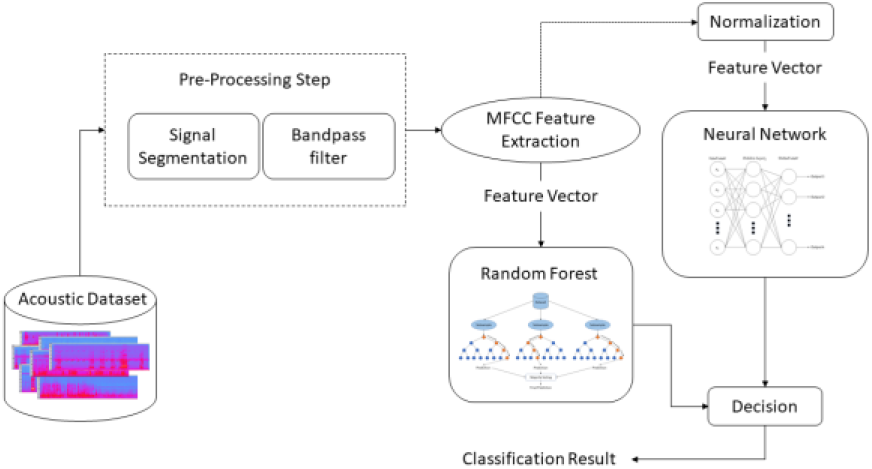
Psittacidae recognition approach.

### 2.2. Datasets

The material used in this paper was collected from 3 different sources containing real-world data. Most of the recordings used to train and validate the machine learning models were taken from the Wikiaves (WikiAves, 2018) and Xeno-canto (Xeno-Canto, 2018) collections. Wikiaves is a community driven website for sharing information, photos and recordings of Brazilian birds. Similarly, Xeno-canto is a website for sharing bird vocalizations from species around the world.

In real world scenarios, it is common for the acoustic landscape to be chaotic, where acoustic events may overlap, vocalizations may be faint, background noise louder, etc. Therefore, datasets that represent this variety of situations are necessary to build a robust method. The Wikiaves and Xeno-Canto collections provided us with this type of data as recordings were collected by means of different microphones, different hardware, varying degrees of background noise, different operators, etc. Furthermore, it is worth noting that as there was a reasonable amount of redundancy between these 2 datasets, we took the precaution of manually analyzing each recording to avoid this issue.

As the objective of this paper is to build a machine learning method to identify particular species among a collection of vocalizations from the same family, we selected 5 other Pscitacidae species that share the same ecosystem and present similar vocalization. Table 1 lists the common name, scientific name and family of all species used in this work.

**Table 1.** BIRD SPECIES

| Scientific Name | Common Name | Family |
| --- | --- | --- |
| *Amazona aestiva* | Turquoise-fronted Amazon | Psittacidae |
| *Amazona amazonica* | Orange-winged Amazon | Psittacidae |
| *Amazona brasiliensis* | Red-tailed Amazon | Psittacidae |
| *Amazona farinosa* | Southern mealy Amazon | Psittacidae |
| *Amazona vinacea* | Vinaceous-breasted Amazon | Psittacidae |
| *Aratinga auricapillus* | Golden-capped Parakeet | Psittacidae |
| *Primolius maracana* | Blue-winged Macaw | Psittacidae |

In total, we collected 196 different recordings of which 64 were from the Xeno-canto database, and 132 from the Wikiaves dataset. These 196 recordings can be divided into 7 classes, according to the subspecies of Psittacidae. In this way, 10 recordings were of the *Amazona aestiva* (AA), 21 classified as *Amazona amazonica* (AM), 64 identified as *Amazona brasiliensis* (AB), and 24 of *Amazona farinosa* (AF), 50 from *Amazona vinacea* (AV), 12 from *Aratinga auricapillus* (AU), and 15 recordings identified as *Primolius maracana* (PM). It is worth noting that, as most of these recordings are over long periods of time and rich in acoustic vocalizations, the segmentation stage will yield multiple examples from each recording for training the neural network.

### 2.3. Pre-processing

Figure 2 shows an example of the vocalization from each Psittacidae species used in this work. As these species often share the same ecosystem, and present vocalizations in similar frequency bands, their acoustic similarity can be easily observed.

**Figure 2.**
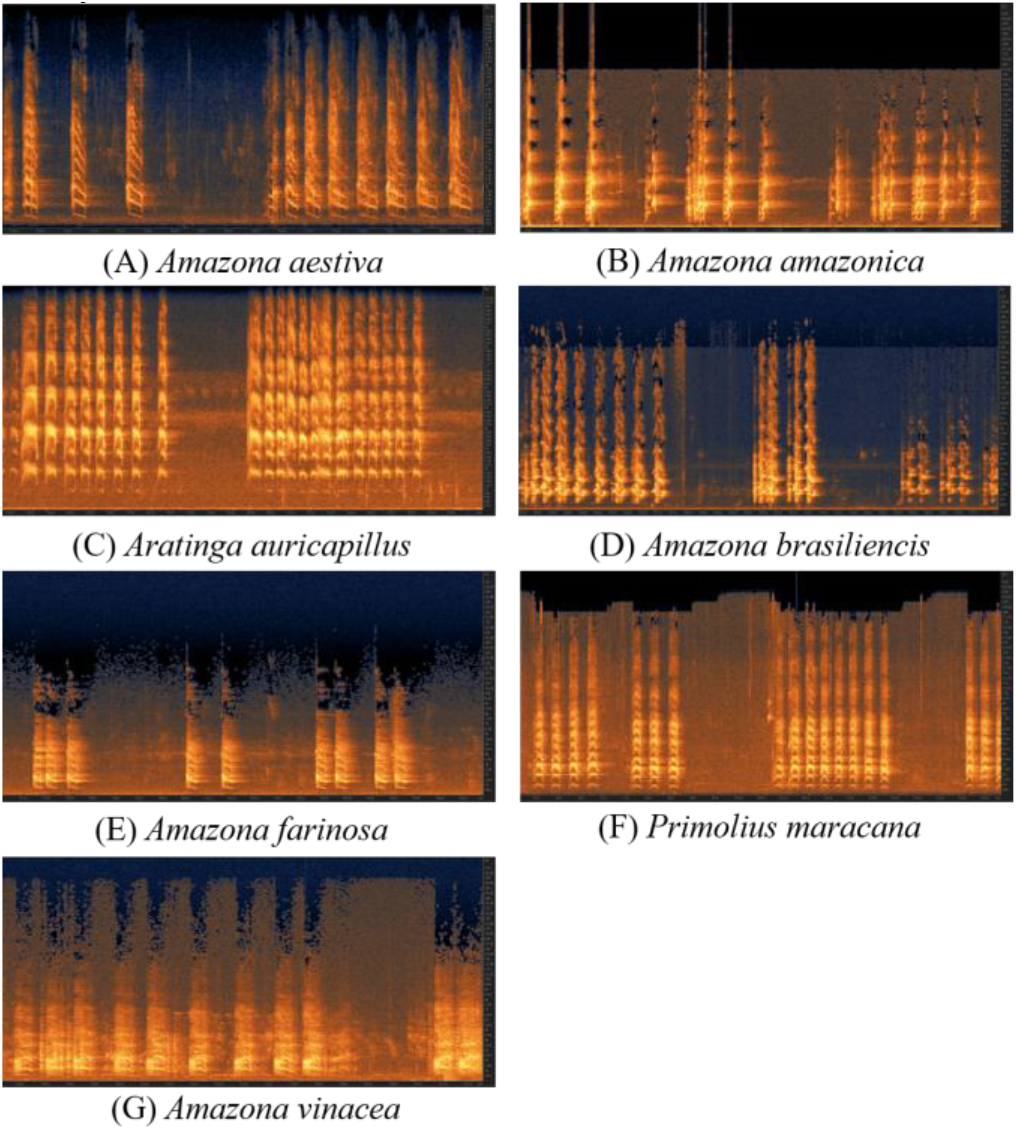
Vocalization samples from all studied Psittacidae species

In addition, as can be better observed from Figure 3 and Figure 4, which shows the vocalization for the *Amazona brasiliensis* and *Amazona vinacea* respectively, recordings normally have a wide range of acoustic events besides the desired signal. Oftentimes, many of these events are not relevant to the problem, such as the vocalization of another animal, background noise such as crickets and instrumentation handling, and may in fact induce the model to learn erroneous features. Therefore, we segmented each recording into smaller time-series containing only the desired signal for the training stage. This segmentation process was performed using the Audacity (Audacity, 1999-2018) software and segmentation tools. Additionally, it is worth noting that occasionally vocalizations overlapped, and in these cases, we selected the segments where the desired signal was still clear to identify.

**Figure 3.**
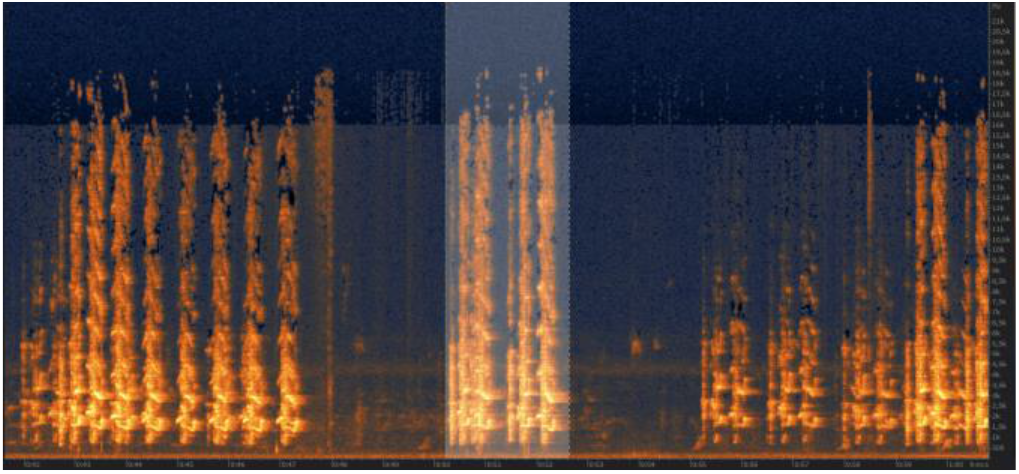
Spectrogram with the vocalization of the *Amazona brasiliensis*

**Figure 4.**
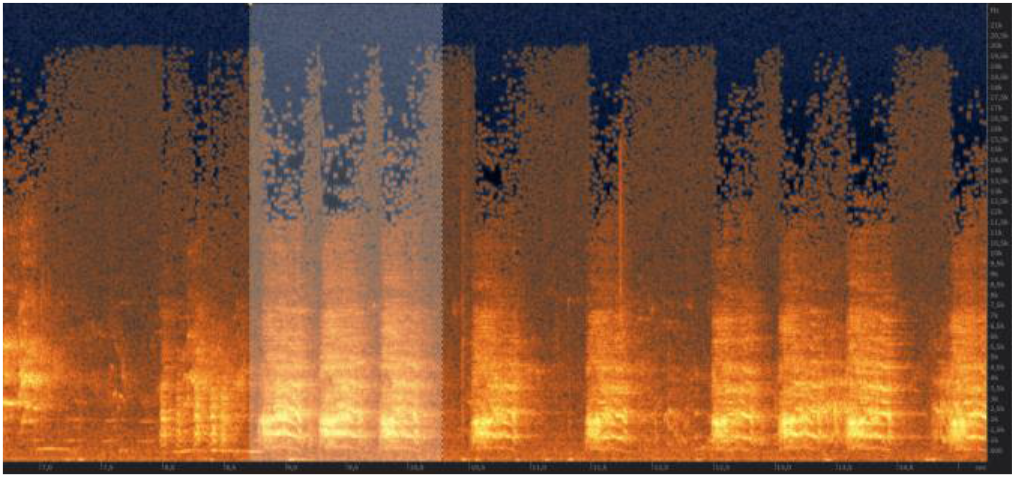
Spectrogram with the vocalization of the *Amazona vinacea*

As can be observed in Figure 2c background noise such as crickets are present across most of the recordings and are detrimental to the classification task. Therefore, in order to reduce interference from background, all recordings were submitted to a bandpass filter to select only the frequencies belonging to the *Amazona brasiliensis* and *Amazona vinacea* vocalization. A bandpass filter is a filter that emphasizes the data in a frequency band of a time-series and attenuates frequencies below or above (Christiano and Fitzgerald, 2003). However, it is worth noting that the choice of an ideal band for filtering a time-series should not be underestimated, as important acoustic features could be lost in the process. Hence, even though ideally, it is preferable to avoid filtering, a trade-off must be weighted, between effectively reducing interference from noise sources and conserving relevant information to the recognition problem.

In this work, after closely analyzing vocalizations of the *Amazona brasiliensis* and *Amazona vinacea* from the collected samples, we conducted a series of preliminary experiments in order to select a reasonable band for the bandpass filter. This experiment consisted in applying a couple of different frequency bands to the samples in order to gauge the best trade-off between noise elimination and information conservation. Table 2 shows the Accuracy for this preliminary experiment with some of the tested bands.

**Table 2.**
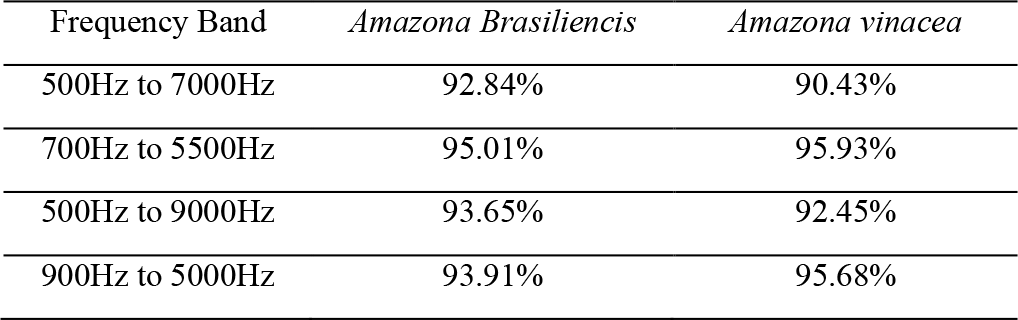
PRELIMINARY EXPERIMENT: ACCURACY

As expected, by analyzing the results of this preliminary experiment, it is possible to identify some promising results as filtering the data in the band 700Hz to 5500Hz improved the classification results. Therefore, for the remainder of this work, all recordings were filtered using a band pass filter between 700Hz and 5500Hz.

### 2.4. MFCC Feature Extraction

The Mel-Frequency Cepstrum coefficients (MFCC) (Foote, 1997) is one of the most widely used methods for extracting features from acoustic events, with applications in both speaker and acoustic species identification (Pace, 2008; Jaffar and Ramli, 2013; 2009; Lopes et al., 2011; Brandes, 2008; Johnson et al., 2002; Selin et al., 2006). Originally developed for application in speech and speaker identification, it is a robust and flexible method, reliable in applications where a lot of outside noise can be expected (Jaffar and Ramli, 2013), making it a simple choice for most scenarios where sound is recorded in an outside environment.

Mel frequency scale relates the perceived pitch, or frequency of a tone, to the real measured frequency, in a way that closely resembles the human auditory system. Basically, humans are better at perceiving variations in a signal at lower frequencies and the mel-scale tries to capture such differences. In its turn, cepstrum is a way to measure the rate of change in the spectral bands and is useful to observe periodic elements (such as peaks in the corresponding frequency spectrum). It is calculated by taking the Fourier transform of a signal into the frequency domain, then taking the logarithm of the magnitude of the resulting frequency spectrum and finally taking the inverse Fourier transform.

The Mel Frequency Cepstrum Coefficients computes the coefficients by a series of filters called the Mel filter bank, which is a series of overlapping triangular shaped bandpass filters distributed on the Mel scale. Then, the log magnitude squared of each mel frequency is computed and its discrete cosine transform calculated. The resulting feature matrix are the MFCC. Moreover, since each sample signal may generate MFCC matrixes with different lengths, they were flattened and zero padded as to have the same dimensions. A more detailed definition of MFCC can be found in (Sigurdsson et al., 2006).

Finally, specifically for the MLP Neural Network, the data was normalized in order to eliminate scale factors present between variables of the data. For the MLP, this procedure is especially important in our scenario since each recording was taken under very different circumstance and settings.

### 2.5. Multilayer Perceptron

Since past decades, Artificial Neural Networks have been used to tackle problems in a multitude of areas for good reason, as they usually present excellent results and are flexible enough to accommodate different settings. Examples of use may be found from medical diagnosis (Moein, 2008) to face recognition (Boughrara *et al.*, 2012), prediction models (Fan *et al.*, 2015), and indeed, also in acoustic species identification (Pace, 2008; Lopes *et al.*, 2011; Selin *et al.*, 2006). There are several types of ANN and, in this work, we utilized the classical Multi-Layer Perceptron (MLP) for its simplicity, general use case covering a wide range of applications, and well documented results.

The MLP consists of a series of interconnected neurons, often called nodes, in 3 layered structures. These structures are called the Input Layer, that receives the features from the training set, one or more hidden layers, and an output layer, that is mapped to the class labels. As shown in Figure 5, each node from one layer is connected to all other nodes from adjacent layers through a structure called axons. Each axon is composed of a weight and the output signal from its node.

**Figure 5.**
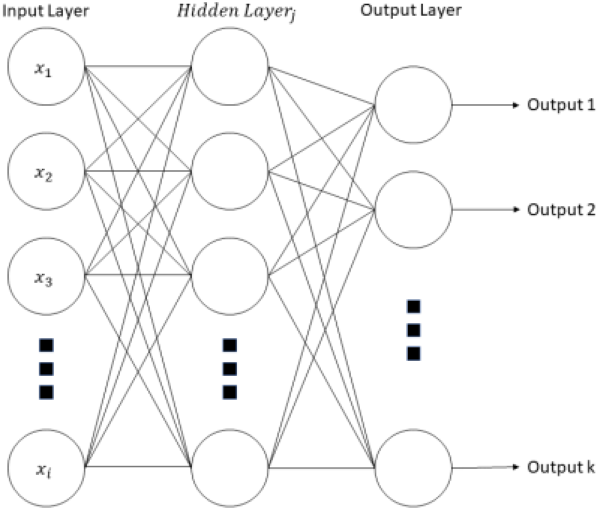
MLP: Each circle represents a unit in its respective layer, and is connected to adjacent units by axons, represented by lines. *x* is the input vector.

Basically, as with other supervised machine learning methods, the MLP learns through a training process by mapping the input features to an output label. After one iteration of the algorithm, the error is calculated by finding the difference between the obtained output and the correct label. Next, during the training stage, the weights of the axons are adjusted in order to minimize the error and arrive as close to the correct answer as possible. This optimization procedure is done through an optimization algorithm, such as the ADAM method (Kingma and Ba, 2014). In this context, the most widely adopted algorithm for the training process is the backpropagation algorithm (Bishop, 1995).

### 2.6. Random Forest Algorithm

The Random Forest algorithm is an ensemble method that trains multiple instances of decision trees on subsets of the data, and then combines them resulting in a more effective and accurate prediction. Figure 6 displays a schematic representation of the Random Forest Algorithm.

**Figure 6.**
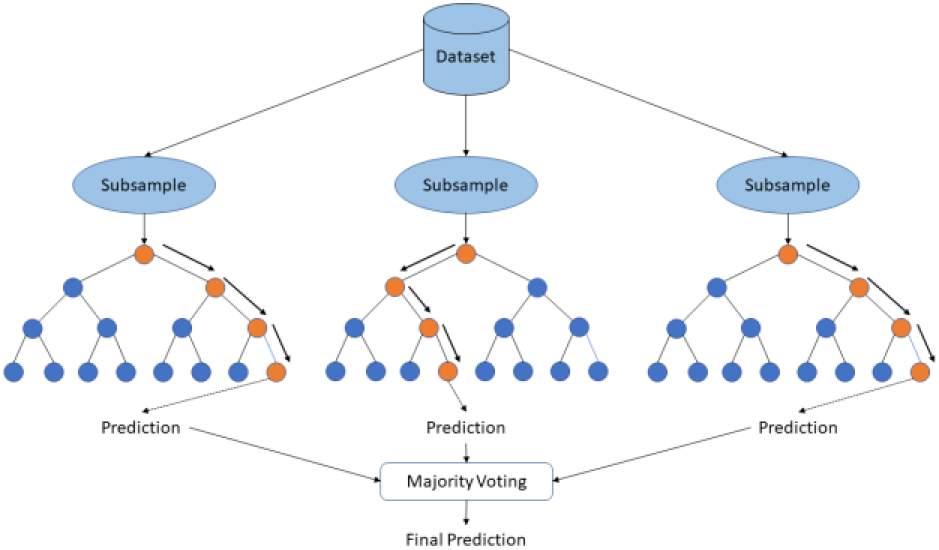
Random Forest Algorithm: Dataset is split into subsamples, and each tree-like structure gives its own prediction. Finally, a majority voting is conducted to arrive at the final prediction.

In general, decision trees (Breiman *et al*, 1984) are a top-down tree-like structure in which each internal node and branches represents conjectures of features (questions) leading up to leaf nodes and each leaf node represents a class label. Classification is performed by applying a majority vote to each leaf node. One drawback of decision trees is that it is very susceptible to small changes in the training data, in what is called high-variance. Ways to mitigate the problem is to use techniques such as ensemble and bagging.

Ensemble methods are based on the general idea that, by combining several weaker instances of a learning algorithm, it is possible to improve the performance of prediction tasks. An initial ensemble method for classification trees, called Random Decision Forests, was proposed by Tin Kam Ho (Ho, 1995). In it, multiple decision trees are trained on a random subspace of the available data. Later, Breiman (Breiman, 2001) proposed the Random Forest algorithm by including an additional bootstrapping aggregation technique, called bagging, that involves randomly sampling a population for each model (that may overlap), from one dataset.

The Random Forest algorithm is one of the most popular machine learning algorithms and has been implemented in a multitude of problems (Khalilia *et al.*, 2011; Rong *et al.*, 2009). It is an easy to use method, capable of presenting great results even without hyper-parameter tuning and can be used for both classification and regression problems. Indeed, across the literature, there are several works that show the effectiveness of the Random Forest algorithm in animal vocalization identification as well (Bowman *et al.*, 2019; Lytle *et al.*, 2010).

## 3. Results and discussions

### 3.1. Experimental Protocol

As described in Section 2.2, all recordings were submitted to a pre-processing step, consisting of signal segmentation and filtering with a bandpass filter between 700Hz and 5500Hz, resulting in a series of acoustic samples. Next, 20 MFCC features were extracted for every frame from each sample, zero-padded to the maximum length from all samples and normalized only for the MLP algorithm. Experiments with more than 20 MFCC features resulted in negligible improvements to justify the additional computational overhead. In this way, the original 196 recordings were converted into 1958 samples that were used to train the Random Forest and MLP classifiers. Of these 1958, 176 were from *Amazona aestiva*, 179 from *Amazona amazonica*, 504 from the *Amazona brasiliensis*, 191 from the *Amazona farinosa*, 535 from *Amazona vinacea*, 179 from *Aratinga auricapillus* and 194 samples from the *Primolius maracana*.

For the MLP algorithm, all experiments were conducted using the ADAM optimization algorithm as it presented good results and is widely used in other works. Preliminary tests with the l-BFGS optimization algorithm showed nearly identical results, on the other hand, the processing time required to train the network increased by a lot when compared to the ADAM method.

In order to evaluate the results obtained from the classification procedure, we used 4 evaluation metrics: accuracy, precision, recall, and the F1 score, described below. Let TP be the number of true positives, TN be the number of true negatives and FP and FN be the number of false positives and false negatives respectively, accuracy can be described as how close the obtained result is to the actual value. Precision tells how many of the selected objects were correct, while recall gives a measure of how many of the correct items were actually selected. Finally, the F1 score is a function of the precision and recall, giving a balance of both. These 4 metrics can be defined as:

- Accuracy: (TP + TN)/(TP+TN+FN+FP)
- Precision: (TP)/(TP+FP)
- Recall: (TP)/(TP+FN)
- 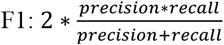

The propose approach was conducted using a machine learning library, the scikit-learn (Scikit-learn, 2011) implementation of the MLP, Random Forest and evaluation methods. All experiments were conducted using a 5-fold stratified cross validation step, where the samples are randomly split 5 times while maintaining class proportion. Next, for each of the folds, the model was trained on the remaining data and evaluated with the one left out. This process produces 5 different results which were then averaged to establish a single final result for each experiment.

### 3.2. Experimental Results

We conducted a series of experiments to evaluate the model against different scenarios, considering the classes (species of Pscitacidae) in our dataset. Firstly, we evaluated the model capability to correctly identify the *Amazona brasiliensis* and *Amazona vinacea* under a complex environment, where all species may be present, that is, *Amazona brasiliensis* vs all other classes (ALL) and *Amazona vinacea* vs ALL (Briggs et al, 2012). Next, we looked at binary classification tasks where different combinations of the classes were tested. For the identification of the *Amazona brasiliensis*, these were AB vs AA, AB vs AM, AB vs AF, AB vs AU, AB vs PM and AB vs AV. For the recognition of the *Amazona vinacea*, the experiments were AV vs AA, AV vs AM, AV vs AF, AV vs AU and AV vs PM. Finally, it is worth mentioning that, for some experiments, we undersampled some classes to combat class imbalance and bias towards the class with most samples.

Tables 3 and 4 present the classification results of the MLP and Random Forest algorithms respectively, under the proposed scenarios.

**Table 3.** PERFORMANCE OF THE NEURAL NETOWORK

|  | Accuracy | Precision | Recall | F1 Score |
| --- | --- | --- | --- | --- |
| AB vs ALL | 81.32% | 83.90% | 81.32% | 80.64% |
| AB vs AA | 97.07% | 97.12% | 97.07% | 97.07% |
| AB vs AM | 93.14% | 93.34% | 93.14% | 93.12% |
| AB vs AF | 88.76% | 89.37% | 88.76% | 88.70% |
| AB vs AU | **98.68%** | **98.73%** | **98.68%** | **98.68%** |
| AB vs PM | 93.14% | 93.40% | 93.14% | 93.12% |
| AB vs AV | 93.94% | 94.17% | 93.94% | 93.92% |
| AV vs ALL | 78.14% | 82.55% | 78.14% | 77.17% |
| AV vs AA | 92.55% | 92.86% | 92.55% | 92.54% |
| AV vs AM | 85.48% | 85.73% | 85.48% | 85.48% |
| AV vs AF | 90.28% | 90.96% | 90.28% | 90.23% |
| AV vs AU | **99.74%** | **99.74%** | **99.74%** | **99.74%** |
| AV vs PM | 93.64% | 93.99% | 93.64% | 93.61% |

**Table 4.**
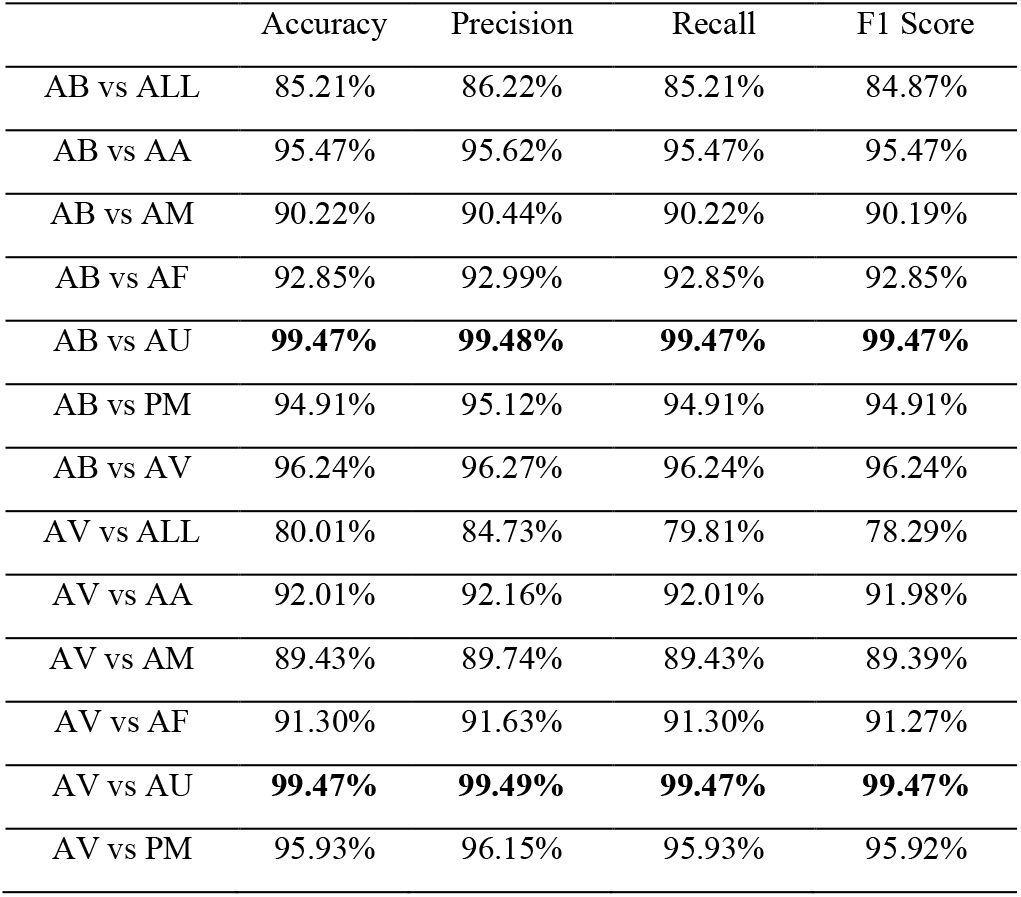
PERFORMANCE OF THE RANDOM FOREST ALGORITHM

|  | Accuracy | Precision | Recall | F1 Score |
| --- | --- | --- | --- | --- |
| AB vs ALL | 85.21% | 86.22% | 85.21% | 84.87% |
| AB vs AA | 95.47% | 95.62% | 95.47% | 95.47% |
| AB vs AM | 90.22% | 90.44% | 90.22% | 90.19% |
| AB vs AF | 92.85% | 92.99% | 92.85% | 92.85% |
| AB vs AU | **99.47%** | **99.48%** | **99.47%** | **99.47%** |
| AB vs PM | 94.91% | 95.12% | 94.91% | 94.91% |
| AB vs AV | 96.24% | 96.27% | 96.24% | 96.24% |
| AV vs ALL | 80.01% | 84.73% | 79.81% | 78.29% |
| AV vs AA | 92.01% | 92.16% | 92.01% | 91.98% |
| AV vs AM | 89.43% | 89.74% | 89.43% | 89.39% |
| AV vs AF | 91.30% | 91.63% | 91.30% | 91.27% |
| AV vs AU | **99.47%** | **99.49%** | **99.47%** | **99.47%** |
| AV vs PM | 95.93% | 96.15% | 95.93% | 95.92% |

The models presented excellent results under most scenarios, with up to 99% accuracy with both MLP and Random Forest. It is worth noting that even the worst scenario achieved classification results of approximately 80% accuracy. Additionally, all evaluation metrics presented similar results in all experiments, showing that the model is robust under different circumstances. Particularly for our 2 target species, AB vs AV, the model was able to clearly differentiate between them, with 96% accuracy for the Random Forest classifier. However, it is possible to see that even though the MLP presented the best results in a couple of scenarios such as AB vs AA and AB vs AM, it showed more fluctuation in its performance, as can be observed from the results obtained in AB vs ALL and AB vs AF for example. In contrast, the Random Forest algorithm displays considerably better results in those experiments, and more consistent results throughout.

In addition to the results shown in tables 3 and 4, we breakdown the outcomes into two confusion matrixes in tables 5 and 6.

**Table 5.** MLP CONFUSION MATRIX BREAKDOWN

|  | TP | TN | FP | FN |
| --- | --- | --- | --- | --- |
| AB vs ALL | 450 | 386 | 156 | 54 |
| AB vs AA | 196 | 168 | 8 | 4 |
| AB vs AM | 188 | 155 | 24 | 12 |
| AB vs AF | 180 | 167 | 24 | 20 |
| AB vs AU | 200 | 176 | 3 | 0 |
| AB vs PM | 188 | 184 | 10 | 12 |
| AB vs AV | 460 | 513 | 22 | 44 |
| AV vs ALL | 505 | 347 | 193 | 30 |
| AV vs AA | 182 | 164 | 12 | 18 |
| AV vs AM | 180 | 144 | 35 | 20 |
| AV vs AF | 183 | 167 | 24 | 17 |
| AV vs AU | 200 | 178 | 1 | 0 |
| AV vs PM | 192 | 182 | 12 | 8 |

**Table 6.**
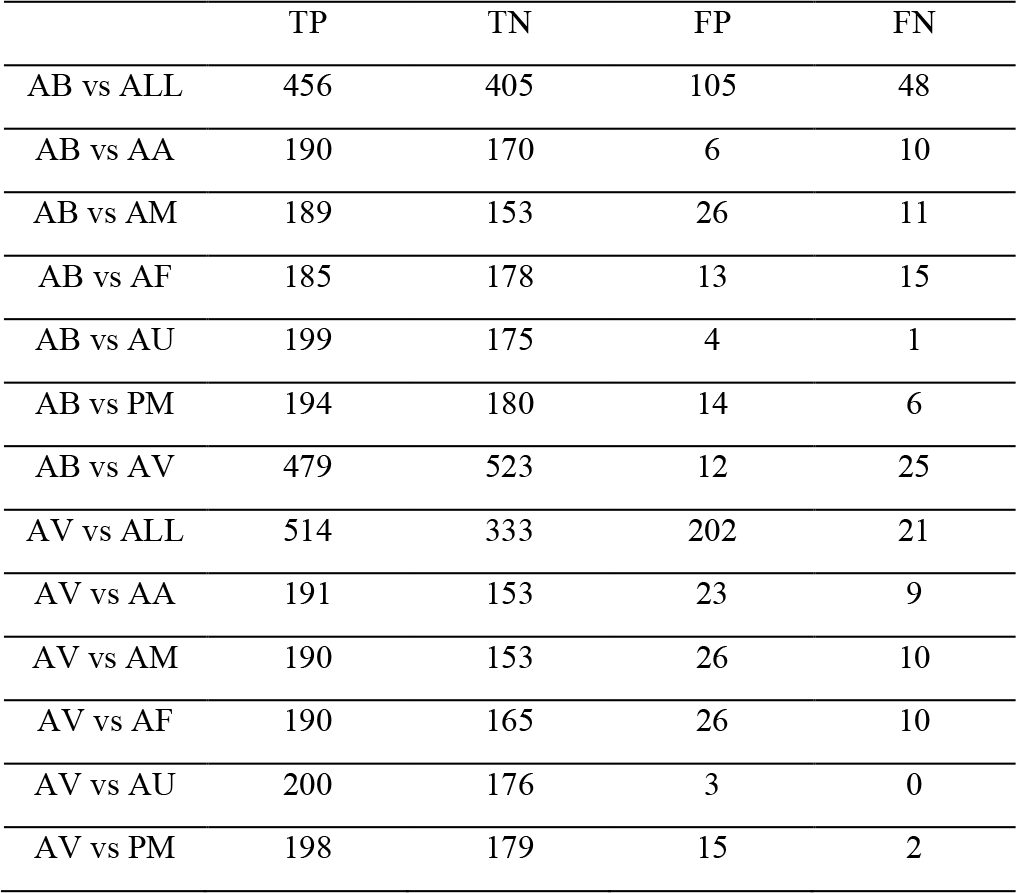
RANDOM FOREST CONFUSION MATRIX BREAKDOWN

It is possible to see that overall, the numbers of false positives in every experiment were higher than the number of false negatives, particularly for the experiments that had lower accuracy such as AB vs ALL and AV vs ALL for both classifiers. We attribute this to the fact that, even though we took some precautions to combat class imbalance, we were able to collect considerably more samples from our target species. For example, in both AB vs ALL and AV vs ALL, the classifiers were tasked with a more complex problem and had access to only a few samples from each wrong class. Hence, the model had more difficulties in correctly identifying the class that was not the *Amazona brasiliensis* or *Amazona vinacea*, resulting in more misses. Therefore, we believe that the classifiers could have achieved even better results by increasing the size of our dataset, notably for the Random Forest Algorithm that showed a strong performance with similar sample quantities. On the other hand, acquiring good field recordings of many different species is not a trivial task, as mentioned in Section 1.

## 4. Conclusion

This work provided an approach for the recognition and identification of 2 endangered parrot species endemic to Brazil, the *Amazona brasiliensis* and *Amazona vinacea,* that oftentimes share the same ecosystem with other species from the same family with similar vocalization.

Two machine learning methods, an ANN and the Random Forest algorithm were trained to tackle the identification task, and their results compared. The proposed approach achieved excellent classification results by both methods, of up to 99% for the Random Forest Algorithm. Indeed, the described methodology was able of correctly identifying the 2 species among similar vocalizations from other 5 from the same family. Furthermore, even the worst results obtained achieved accuracy of up to 80%. Thus, the method proved to be effective, flexible and robust to be considered for this type of task.

In this context, we contemplate the following as future work: (i) the development of a real time, fully automated PAM module utilizing this approach; (ii) apply the method in a bigger real soundscape database, that is currently being constructed by continuous recording from multiple sensors; (iii) implement this method in conservation units in Brazil.

